# A strategic sampling design revealed the local genetic structure of cold-water fluvial sculpin: a focus on groundwater-dependent water temperature heterogeneity

**DOI:** 10.1101/2021.03.19.435938

**Authors:** Souta Nakajima, Masanao Sueyoshi, Shun K. Hirota, Nobuo Ishiyama, Ayumi Matsuo, Yoshihisa Suyama, Futoshi Nakamura

## Abstract

A key piece of information for ecosystem management is the relationship between the environment and population genetic structure. However, it is difficult to clearly quantify the effects of environmental factors on genetic differentiation because of spatial autocorrelation and analytical problems. In this study, we focused on stream ecosystems and the environmental heterogeneity caused by groundwater and constructed a sampling design in which geographic distance and environmental differences are not correlated. Using multiplexed ISSR genotyping by sequencing (MIG-seq) method, a fine-scale population genetics study was conducted in fluvial sculpin *Cottus nozawae*, for which summer water temperature is the determinant factor in distribution and survival. There was a clear genetic structure in the watershed. Although a significant isolation-by-distance pattern was detected in the watershed, there was no association between genetic differentiation and water temperature. Instead, asymmetric gene flow from relatively low-temperature streams to high-temperature streams was detected, indicating the importance of low-temperature streams and continuous habitats. The groundwater-focused sampling strategy yielded unexpected results and provided important insights for conservation.

## 1. Introduction

How genetic structure is shaped across a landscape is an essential theme in evolutionary biology, and this information provides an empirical basis for conservation biology by elucidating habitat connectivity or predicting the effects of landscape change (Hohenlohe et al. 2021). A pattern of ‘isolation by distance’ (IBD; Wright 1943), in which genetic differentiation increases with geographic distance, is a very common pattern to explain population structure (Meirmans 2012). In addition to space, another important component of a landscape is the environment (Nosil et al. 2005; Thorpe et al. 2008). In the field of landscape genetics, many studies have investigated the interactions between genetic differentiation and landscape variables, and a couple of patterns have described the relationship between genetic structure and environment. When gene flow between populations that inhabit different environments is limited due to local adaptation or other factors, genetic differentiation increases with environmental differences and a pattern of ‘isolation by environment’ (IBE; Wang and Summers 2010) is generated. Another pattern that involves genetic structure and the environment is ‘counter-gradient gene flow’ (Sexton et al. 2014). In this scenario, gene exchange between different environments is high and forms a source-sink pattern in a landscape. Because these interactions between genetic differentiation and landscape variables are critically important for addressing classical evolutionary questions related to ecological speciation or conservation, the environment is a factor that cannot be ignored when describing the observed population structure (Orsini et al. 2013; Wang and Bradburd 2014).

However, it is very difficult to disentangle the relative strength of space (distance) and environment in observed patterns of spatial genetic differentiation because environmental differences are usually highly correlated with geographic distance (Sexton et al. 2014; Wang and Bradburd 2014). To test spatial relationships, the Mantel test and partial Mantel test are commonly used. However, the Mantel test family is strongly criticized for its inflated Type I error rate, especially under conditions where the measured variables are spatially correlated (Guillot and Rousset 2013; Harmon and Glor 2010; Meirmans 2012; Raufaste and Francois 2001; Rousset 2002). Although there are several proposed alternative methods to the Mantel test such as multiple regression on distance matrices (MRM; Lichstein 2007), Procrustes analysis, or redundancy analysis (RDA), and some of them perform much better than the partial Mantel test, previous studies have suggested that all methods fail to correctly model the relative importance of space and environment under certain patterns, which cannot be known *a priori* (Diniz-Filho et al. 2013; Gilbert and Bennett 2010). It is now recognized that no method has the ability to address the spatial-environmental correlation and perform hypothesis testing simultaneously (Zeller et al. 2016). These studies suggest the difficulty of separating spatial and environmental effects on genetic differences. However, once a sampling design that decouples distance and environment is developed, we will be able to clearly compare the relative strength of space and environment independent of these controversies. In the field of community ecology, Gilbert and Lechowicz (2004) used a sampling design that removed spatial autocorrelation of the environment sampled, and they analyzed the species’ spatial and environmental correlations to reject the neutral theory of biodiversity. However, subsequent attempts to partition spatial and environmental components have focused exclusively on statistical and analytical methodology (Gilbert and Bennett 2010). Especially in landscape genetics, sampling design has not received much attention (Meirmans 2015).

Here, we focused on stream ecosystems, which often exhibit heterogeneous environments (Heino et al. 2013; Uno 2016). In stream ecosystems, water temperature is the strongest factor affecting river organisms as well as flow regime (Olden and Naiman 2010), and it is increasingly being an important variable under ongoing climate change. Water temperature is determined by air temperature, riparian forests, and groundwater, among others (Caissie 2006; Nakamura and Yamada 2005). In particular, the effect of groundwater discharge on water temperature has been indicated to be very strong (Arscott et al. 2001). However, the influence of groundwater discharge on the upper sections of streams has received much less attention (Brown et al. 2007), and many ecological studies have used air temperature instead of water temperature (e.g., Almodóvar et al. 2012; Middaugh et al. 2018). Groundwater shows a very stable water temperature and flow regime; hence, groundwater discharge greatly affects the stream environment (Poff et al. 2010). Given water temperature in summer that is critical for cold-water organisms, streams with high groundwater display lower values and these streams could be an important habitat for these species. Since the amount of cold groundwater discharge vary locally within a watershed, groundwater creates spatial heterogeneity of environments within the watershed. Thus, to reveal the ecological role of water temperature, knowledge of the spatial heterogeneity caused by groundwater is thought to be useful.

As a practical study, we investigated the genetic structure of the cold-water fluvial sculpin *Cottus nozawae* in the upstream section of the Sorachi River, Hokkaido, Japan. Cold-water fish are very vulnerable to climate change, and it is worth studying the relationships between population structure and water temperature. Together with *C. nozawae*, *Salvelinus malma*, *Salvelinus leucomaenis*, *Barbatula oreas*, and *Parahucho perryi* inhabit this watershed. However, studies on *S. malma* and *S. leucomaenis* did not display strong genetic structure within this watershed due to their active migration (Koizumi 2011; Nakajima et al. 2020). Due to the lower mobility of adult *C. nozawae* than these species (Goto 1998; Okumura and Goto 1996), we predicted that *C. nozawae* should display clear genetic differentiation within the watershed. In addition, the summer water temperature has been shown to be the dominant factor determining the distribution and survival of this species (Yagami and Goto 2000), and a relationship in which low water temperature in summer increases the density of this species has also been confirmed in our study area (Suzuki, unpublished). Therefore, summer water temperature can be explicitly used as an important variable for *C. nozawae*. Hypothesis-driven studies that focus on a given variable have advantages over exploratory studies in designing sampling strategies (Richardson et al. 2016). Since ecologically important variables affecting survival and population density could cause local adaptation between different environments (Kawecki and Ebert 2004), we predicted that the differences in summer water temperature cause local adaptations and lead to the IBE pattern within the watershed.

The aims of this study are (i) to eliminate the correlation of geographic distance and water temperature differences via a groundwater-focused sampling strategy, (ii) to investigate the local genetic structure of *C. nozawae* and its determinant factors, and (iii) to discuss the relationship between water temperature and the population structure of cold-water fish. Although recent studies have frequently used adaptative genetic markers to detect local adaptation, barriers to gene flow imposed by selection and local adaptation between populations can be detected with neutral markers (Sexton et al. 2014). Given that the use of adaptive markers is still challenging, the evaluation of current genetic differentiation and connectivity based on neutral genetic markers is still informative and can assist in allocating conservation units to preserve local genetic variation (Tsuda et al. 2015). To assess the genetic structure, we used putatively neutral genome-wide SNPs on the inter simple sequence repeat (ISSR) region and analyzed genetic differentiation and gene flow patterns.

## 2. Materials and Methods

### 2.1 Study sites and sampling

This study was conducted in the upper section of the Sorachi River, Hokkaido, Japan (Fig. 1; Table 1). To conduct sampling across a heterogeneous landscape, we used an approach based on watershed geology, which is known to cause significant variations in water temperature through groundwater discharge (Caissie 2006; Nagasaka and Sugiyama 2010; Tague et al. 2007). In the Sorachi River, many small spring-fed inputs are found in the Quaternary volcanic region (Koizumi and Maekawa 2004; Watz et al. 2019). Thus, the sampling sites were selected so that tributaries with volcanic watersheds and nonvolcanic watersheds were spatially intermingled. We also chose sites that ensured that the watershed was spatially well represented (small spatial gaps). Watershed geology was assessed based on the Seamless Digital Geological Map of Japan V2 from the National Institute of Advanced Industrial Science and Technology. There is a large dam (Kanayama Dam) in the watershed. While the area upstream of Kanayama Dam represents one of the most continuous river habitats in Japan and there are no barriers between populations, the downstream region is slightly more influenced by anthropogenic impacts on streams. For example, one population (Pop13) is located upstream of a check dam, and another population (Pop19) is located in a tributary with a downstream portion that runs through an agricultural landscape.

**Table 1.** Details of the sampling sites and genetic diversity of each population.

| Population ID | Localities | Longitude (E°) | Latitude (N°) | Altitude (m) | N | $H_E$ | $F_{IS}$ |
| --- | --- | --- | --- | --- | --- | --- | --- |
| 1 | Tomamu River | 142.670 | 43.057 | 526 | 32 | 0.121 | 0.016 |
| 2 | Kinnosawa River | 142.672 | 43.085 | 466 | 22 | 0.113 | 0.005 |
| 3 | Shikerebenaizawa River | 142.699 | 43.107 | 451 | 31 | 0.112 | 0.006 |
| 4 | Ehoroakanbetsu River | 142.699 | 43.218 | 510 | 34 | 0.114 | 0.012 |
| 5 | Ozawa River | 142.703 | 43.218 | 511 | 30 | 0.113 | 0.004 |
| 6 | A tributary of the Seesorapuchi River | 142.678 | 43.163 | 445 | 21 | 0.111 | 0.009 |
| 7 | Pankeyara River (up) | 142.699 | 43.155 | 450 | 32 | 0.112 | 0.007 |
| 8 | Pankeyara River (down) | 142.680 | 43.133 | 414 | 32 | 0.114 | 0.001 |
| 9 | Peiyurushiebe River (up) | 142.739 | 43.136 | 498 | 32 | 0.108 | 0.007 |
| 10 | Peiyurushiebe River (down) | 142.680 | 43.129 | 413 | 32 | 0.113 | 0.014 |
| 11 | Ecchudantai-no-sawa River | 142.624 | 43.137 | 387 | 32 | 0.113 | -0.001 |
| 12 | Ikutora River | 142.603 | 43.173 | 376 | 32 | 0.113 | 0.009 |
| 13 | Tonashibetsu River (up) | 142.335 | 43.118 | 378 | 22 | 0.085 | 0.003 |
| 14 | Tonashibetsu River (mid) | 142.346 | 43.135 | 340 | 21 | 0.106 | 0.007 |
| 15 | Tonashibetsu River (down) | 142.393 | 43.149 | 289 | 22 | 0.113 | 0.014 |
| 16 | Nishitappu River (up) | 142.593 | 43.224 | 375 | 22 | 0.115 | 0.003 |
| 17 | Nishitappu River (mid) | 142.558 | 43.207 | 325 | 22 | 0.117 | 0.000 |
| 18 | Nishitappu River (down) | 142.501 | 43.216 | 292 | 19 | 0.112 | -0.001 |
| 19 | Roseppu River | 142.463 | 43.260 | 336 | 21 | 0.101 | 0.002 |
| 20 | Onkosawa River | 142.403 | 43.218 | 239 | 20 | 0.108 | 0.038* |

N, number of individuals; $H_E$ , expected heterozygosity; $F_{IS}$ , fixation index (significant deviation from zero is denoted by asterisks; \* $p < 0.05$ after Bonferroni correction).

**Fig. 1.**
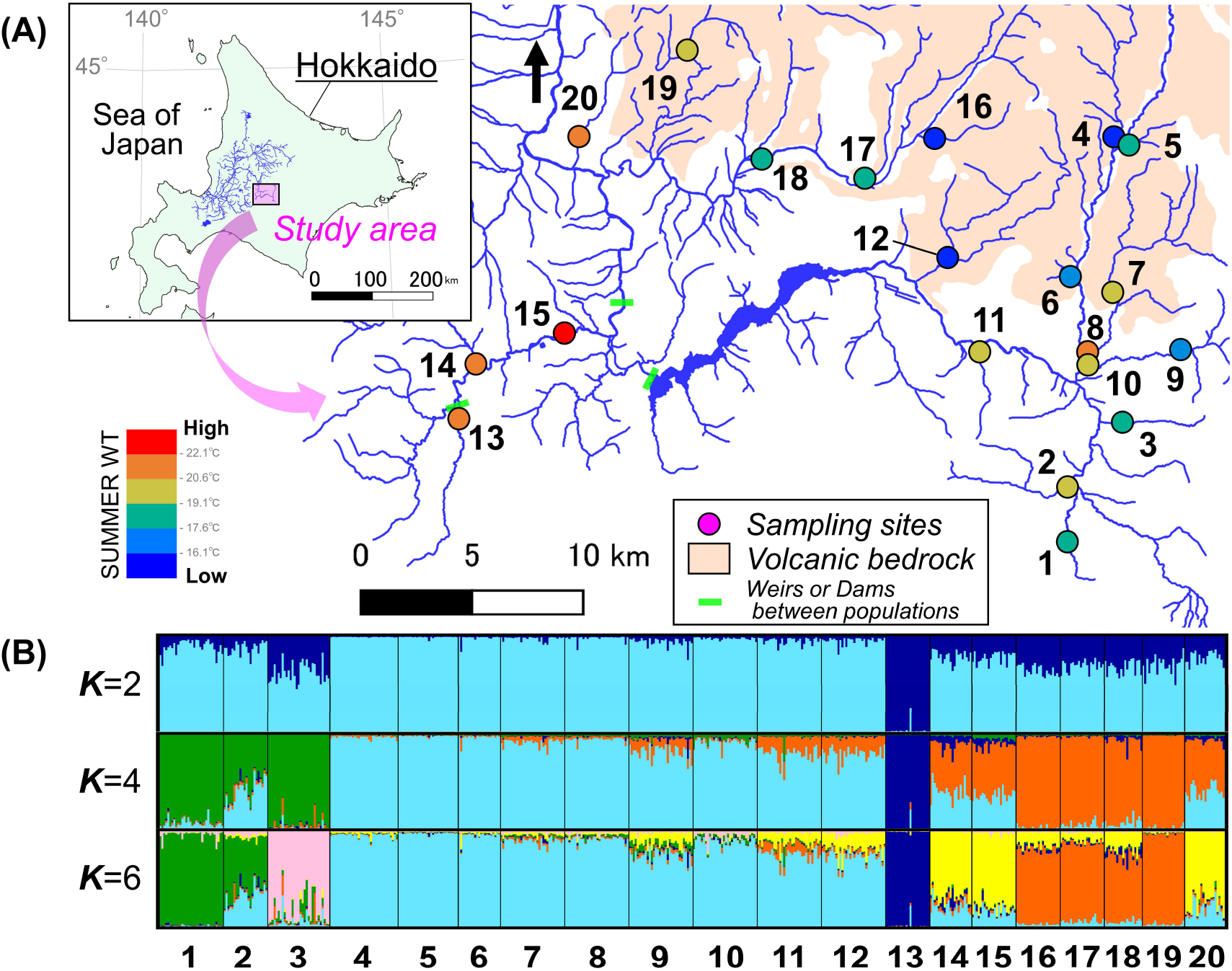
(A) Study area. Fill colors of the sampling site depend on the maximum water temperature. Site labels correspond to population IDs in the text and Table 1. (B) Population structure inferred by STRUCTURE. The number indicates the population IDs.

From 20 sites in the Sorachi River, 531 individuals of *C. nozawae* were caught for genetic analysis. The fish were caught by electrofishing (model 12-B Backpack Electrofisher; Smith-Root Inc.) or with hand nets. Small pieces of fin tissue were clipped, placed in 99.5% ethanol, and stored at −20 °C in the laboratory until DNA extraction. Total DNA was extracted using QIAGEN DNeasy Blood and Tissue Kit (QIAGEN Inc.) with a combination of Genomic DNA Extraction Column (FAVORGEN Inc.).

### 2.2 Water temperature

At each sampling site, the summer water temperature was recorded hourly using data loggers (Onset Computer Corp. or Gemini Data Loggers Ltd., depending on site). Water temperatures were measured during the summer of 2020, and also during the summer of 2019 at some locations. To align the data sampling periods, data from 16 July 2020 to 31 August 2020 were extracted, and the maximum water temperature in this period was identified. At one site (Pop10) where we failed to record water temperatures in some mid-summer periods, the maximum temperature in 2019 was used as an alternative. We used the maximum water temperature which significantly affects the populations of cold-water fish as a representative of summer water temperature (Yagami and Goto 2000). The correlation of water temperature differences with waterway geographic distance (hereafter referred to as geographic distance) was examined with the Mantel test (9999 permutations) using the package ECODIST 2.0.3 (Goslee and Urban 2007) in R 3.6.0 (R Core Team 2019). Then, a Mantel correlogram showing the spatial correlation of water temperature differences across multiple ranges of geographic distance was constructed. The Mantel correlogram in 10 equal distance classes was assessed with 9999 permutations using the package VEGAN 2.5.6 (Oksanen et al. 2019) in R.

### 2.3 MIG-seq library preparation and sequencing

To obtain neutral genome-wide SNP data, we used multiplexed ISSR genotyping by sequencing (MIG-seq; Suyama and Matsuki 2015), a microsatellite-associated DNA sequencing technique. MIG-seq is one of the reduced representation sequencing methods that includes restriction site-associated DNA sequencing (RAD-seq), but the number of available informative loci (MIG-loci) detected by MIG-seq is less than that of RAD-seq, and most of the MIG-loci are putatively neutral. Although MIG-seq is not suitable for outlier analysis or gene identification, acquired loci are sufficient for population genetic analysis. A MIG-seq library was prepared following the protocol outlined in Suyama and Matsuki (2015), except for the minor modifications outlined below. The first PCR was conducted using eight ISSR primer sets with tail sequences at an annealing temperature of 38 °C. The second PCR was conducted using primer pairs including tail sequences, adapter sequences for Illumina sequencing, and five-base (forward) and nine-base (reverse) barcode sequences to identify each individual sample. The conditions for the second PCR were as follows: 12 cycles of denaturation at 98 °C for 10 s, annealing at 54 °C for 15 s, and extension at 68 °C for 1 min. Fragments in the size range of 400–800 bp in the purified library were isolated. After library quantification, the products were sequenced on the Illumina MiSeq platform using MiSeq Reagent Kit v3 following the manufacturer’s protocol.

### 2.4 SNP detection

After removing the primer regions and low-quality reads according to Suyama and Matsuki (2015), a total of 39,956,740 reads were obtained. SNP selection was performed using STACKS 2.41 (Catchen et al. 2013) with the following parameters: minimum depth option creating a stack (*m*) = 3, maximum distance between stacks (*M*) = 2, maximum mismatches between loci when building the catalog (*n*) = 2, and number of mismatches allowed to align secondary reads (*N*) = 4. After the process of *ustacks*, *cstacks*, *sstacks*, *tsu2bam*, and *gstacks* were carried out, the dataset of assembled loci (*stacks dataset*) was obtained. For most analyses (except for section *2.8*), the applied SNP filtering criteria under the *populations* command were as follows: only loci present at a rate of more than 80% among all samples were extracted; the minimum minor allele count was three; sites showing excess heterozygosity (>0.6) were removed; and the output was limited to one SNP per locus. After filtering, 489 SNPs were obtained.

### 2.5 Genetic diversity and differentiation

For each population, the expected heterozygosity (*H*_E_) and fixation index (*F*_IS_) were calculated using the *populations* command in STACKS. Significant deviations from Hardy–Weinberg equilibrium (HWE), as indicated by *F*_IS_ deviating from zero, were tested by 1000 randomizations using FSTAT 2.9.4 (Goudet 1995). Genetic differentiation among *populations* was evaluated by calculating global/pairwise *F*_ST_ values (Weir and Cockerham 1984) using GenAlEx 6.5 (Peakall and Smouse 2012). To understand the spatial trend of the genetic structure, a Mantel correlogram displaying correlations between *F*_ST_ and geographic distance for each of the 10 distance classes was constructed with 9999 permutations using the package VEGAN in R.

### 2.6 Population structure

The population structure was examined using STRUCTURE 2.3.4 (Pritchard et al. 2000), which implements a Bayesian clustering method using multi-locus allele frequency data. The STRUCTURE settings were the admixture and allele frequency correlated model with previous sampling location information (LOCPRIOR; Hubisz et al. 2009). The algorithm was run 20 times for each K from 1 to 14 with a burn-in of 20,000 followed by 30,000 Markov chain Monte Carlo (MCMC) replicates. The program CLUMPAK (Kopelman et al. 2015) was used to compile the results of the STRUCTURE analysis for each K. STRUCTURE HARVESTER (Earl and vonHoldt 2012) was employed to calculate the probability of the data for each K (LnP(D); Pritchard et al. 2000), the corresponding standard deviation, and Evanno’s delta K (Evanno et al. 2005).

### 2.7 Association of genetic variation with geographic distance and water temperature

To evaluate the effects of space and water temperature on genetic differentiation, the independent correlations of *F*_ST_ with water temperature differences and geographic distance were calculated by Mantel tests with 9999 permutations. To identify confounding effects between space and water temperature if present, multiple regression on distance matrices (MRM; Lichstein 2007) was used, with “water temperature differences + geographic distance” as the explanatory variable. Mantel tests and MRM were performed with the package ECODIST in R. These analyses were conducted at three scales: (a) using all 20 populations (waterway distance 0.3-70.6 km), (b) using upstream 12 populations (Pop1-12; waterway distance 0.3-24.1 km), and (c) using structured 9 populations (Pop4-12; populations assigned as one large cluster in STRUCTURE). We used scale (b) to account for the mobility of *Cottus* and to exclude the effects of a long spatial gap and a dam, and scale (c) was used to prevent biases when including sites displaying high genetic divergence (‘outliers’; Koizumi et al. 2006).

Some studies have considered it inappropriate to apply the Mantel test to the raw vector data and to take the difference to create a matrix (Legendre and Legendre 2012). Hence, we also used distance-based RDA, which is a constrained ordination approach that can use environmental variables as vector data and has higher power than the Mantel test (Harmon and Glor 2010; Legendre et al. 2015). We used genetic variation as the response variable and water temperature and spatial variables derived from geographic distance as explanatory variables. For genetic variation, a principal coordinate analysis (PCoA) of the pairwise *F*_ST_ matrix was conducted, and all axes were used as response variables. Spatial predictors were generated as a set of distance-based Moran’s eigenvector maps (MEMs; Griffith and Peres-Neto 2006), vectors that capture broad- to small-scale spatial structures. MEMs were generated from the geographic distance matrix using the package ADESPATIAL 0.3.8 (Dray et al. 2020) in R. To identify meaningful MEM predictors, forward selection (999 permutations) was used for generated MEM dataset. In each RDA with the explanatory variables of water temperature and geographic MEM variables, adjusted R-square values (R_adj_^2^), which penalize the increase in explanatory power due to an increase in the number of explanatory variables, were calculated. RDAs were performed on three scales, as in the Mantel tests and MRM analyses. RDAs including forward selections were performed with the package VEGAN in R.

### 2.8 Contemporary gene flow

To estimate contemporary gene flow rates, BA3SNP (Mussmann et al. 2019), a reconstructed version of BayesAss (Wilson and Rannala 2003) that estimates dual-direction pairwise migration rates over a few generations, was used for the 20 populations. To improve the convergence of BayesAss, another SNP filtering criterion that increases the variance among populations was used. From the *stacks dataset*, loci present at a rate of more than 40% among at least two populations were extracted with a minimum minor allele count of three. Sites showing excess heterozygosity (>0.6) were removed, and the output was limited to one SNP per locus. Then, 1291 SNPs were obtained. We set 20,000,000 MCMC iterations, including a burn-in period of 10,000,000. Following the proposal of Meirmans (2014), we performed 10 independent runs with different seed values and chose the run with the lowest Bayesian deviance to overcome poor MCMC sampling. The inferred migration rates with 95% credible intervals that did not include zero were regarded as significant. We then defined the *net immigration rate* into a given population A, from another population B, as the [estimated value of B->A minus that of A->B]. Then, the *mean net immigration rate* for each population (the mean net immigration rate for a given population from all of its paired net immigration rate values between all other populations) was calculated to measure the degree to which a population is a donor or a recipient of migrants (Hanfling and Weetman 2006; Sexton et al. 2016). To investigate whether water temperature affects the source-sink structure independent of upstream-downstream dispersal, we constructed a linear model with a response variable of the mean net immigration rate and explanatory variables as maximum water temperature and elevation. The Pearson’s correlation between water temperature and elevation was −0.39; thus, no multicollinearity was considered. The independent effects of water temperature and elevation were assessed by the partial regression coefficient (*β*) and standardized partial regression coefficient (std *β*). Likelihood ratio tests were used to determine the significance of explanatory variables. This analysis was conducted using the package LMTEST (Zeileis and Hothorn 2002) in R.

## 3. Results

### 3.1 Heterogeneity of water temperature

Summer water temperatures varied within the watershed, and the maximum water temperature ranged from 12.5 to 23.5 °C. The correlation between geographic distance and water temperature difference was low (Mantel r = 0.08, *p* = 0.45). In the Mantel correlogram for water temperature, r values were not significant in all distance classes (Fig. 2A), indicating little spatial autocorrelation.

**Fig. 2.**
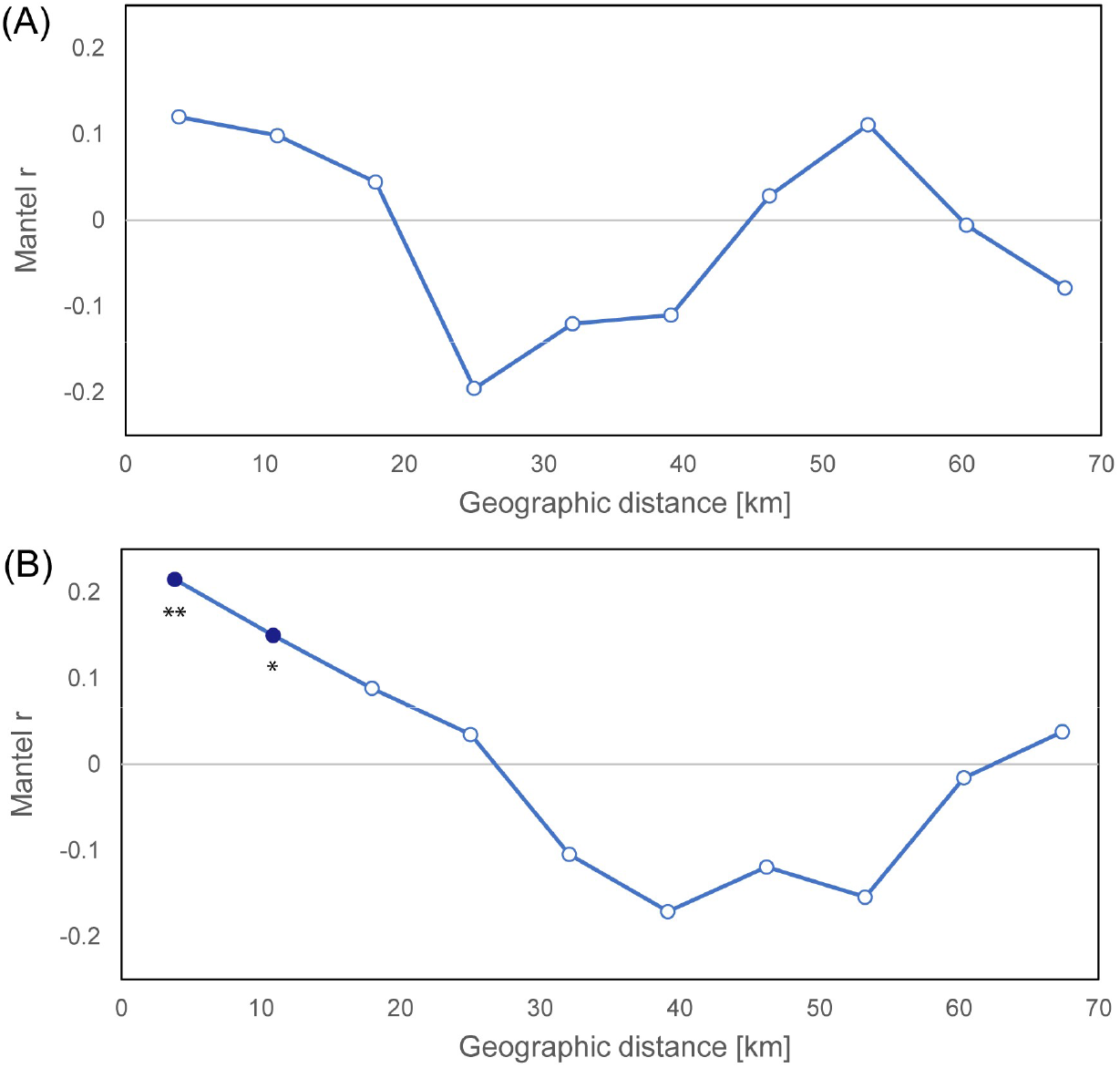
Mantel correlograms presenting spatial autocorrelation of (A) water temperature and (B) genetic data. Filled circles indicate significant correlations from a null model of spatial structure (* *p* < 0.05, ** *p* < 0.01).

### 3.2 Genetic diversity, divergence, and population structure

*F*_IS_ did not deviate significantly from zero except in one population (Pop20), suggesting that HWE could be assumed in most populations. The *H*_E_ was similar across the watershed, but one population located in the upstream section of a check dam (Pop13) displayed a slightly lower value (Table 1). Across the 20 populations, the *F*_ST_ value was 0.038, indicating weak population differentiation, and pairwise *F*_ST_ values ranged from 0.001 to 0.127 (Table S1). In a Mantel correlogram, r values were significantly positive in up to the second distance classes (approximately 15 km) and reached zero at a distance of 27 km (Fig. 2B).

In the STRUCTURE analysis, the probability of the data (LnP(D)) increased progressively with each K, and delta K was highest at K = 4 (Fig. S1). Distinct clusters corresponding to the geographic structure were detected up to K = 6 (Fig. 1). For K = 2, the genomes of the individuals in Pop13 were grouped into a single unique cluster. For K = 4, Pop1-3 and Pop16-19 were grouped as additional clusters. Pop14,15,20 was inferred to be an admixed cluster when K = 4 and a distinct unique cluster when K = 6. Pop3 also formed a unique cluster at K = 6. Pop4-12 was assigned to almost one cluster at even large K.

### 3.3 Effects of space and water temperature on genetic differentiation

Mantel tests captured significant correlations between pairwise *F*_ST_ and waterway geographic distance (Mantel r = 0.27, p < 0.05; Fig. 3), indicating a significant effect of IBD. Conversely, no correlation between *F*_ST_ and water temperature differences was detected (Mantel r = −0.10, p = 0.56). The MRM with these two matrices as an explanatory variable did not show a significant relationship, and the R^2^ value was not very different from that of geographic distance alone. The same pattern was obtained at different spatial scales (Table 2). The forward selection of the MEM variables identified six significant predictors and the RDA with selective MEMs was significant with an R_adj_^2^ value of 0.61 (Fig. S2). Even in RDA, water temperature did not explain the genetic variation at all.

**Table 2.** Summary of Mantel tests, multiple regression on distance matrices (MRM), and RDAs on three spatial scales for examining the effect of space and water temperature on genetic differentiation (*F*_ST_).

| Explanatory matrices | All<br>20 populations | Upstream<br>12 populations | Structured<br>9 populations |
| --- | --- | --- | --- |
| Mantel test ($r^2$) | | | |
| Water temperature | 0.010<sup>†</sup> ($p=0.56$) | 0.023<sup>†</sup> ($p=0.53$) | 0.000 ($p=0.94$) |
| Geographic distance | 0.072 ($p<0.05$) | 0.074 ($p=0.24$) | 0.298 ($p<0.05$) |
| MRM ($R^2$) | | | |
| Water temperature<br>+ Geographic distance | 0.087 ($p=0.13$) | 0.116 ($p=0.32$) | 0.304 ($p<0.05$) |
| RDA ($R_{adj}^2$) | | | |
| Water temperature | -0.017 ($p=0.51$) | -0.073 ($p=0.92$) | -0.083 ($p=0.69$) |
| Geographic distance<br>(MEMs) | 0.609 ($p<0.01$) | 0.374 ($p<0.001$) | 0.486 ($p<0.05$) |

Mantel $r$ values were squared to compare MRM $R^2$ values. $R_{adj}^2$ values can take negative values when the relation is very small. $p$ -values were obtained by two-tailed tests. Upstream 12 populations, Pop1-12; Structured 9 populations, Pop4-12. <sup>†</sup>: original $r$ values were negative.

**Fig. 3.**
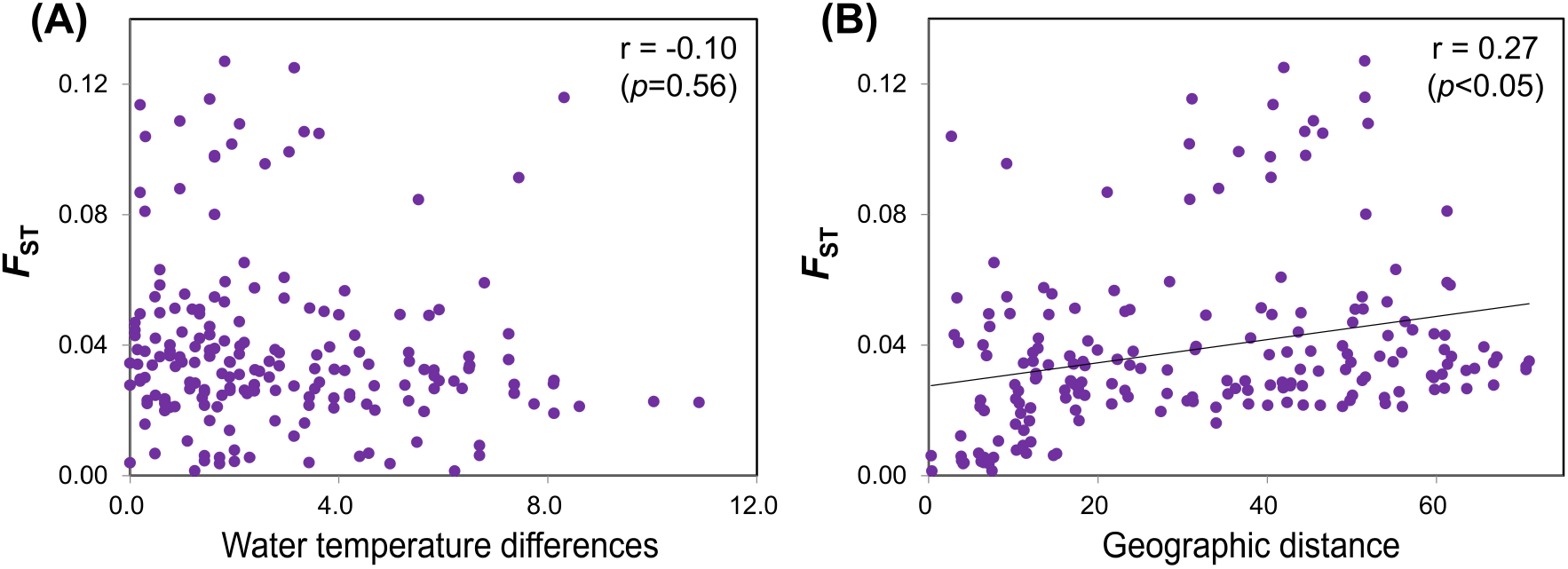
Relationship of genetic differentiation with (A) water temperature and (B) geographic distance.

### 3.4 Gene flow

A total of 11 significant gene flow were detected (Tables 3 and S2). All of them were gene flow from relatively low-temperature sites (mean 14.3 °C [sd 2.90]) to high-temperature sites (mean 20.0 °C [sd 1.84]). Most of them were upstream to downstream direction, but they also included gene flow from downstream to upstream direction (Pop4 to 5 and Pop12 to 11). The linear model indicated that water temperature had a significant effect on the mean net immigration rates (std *β* = 0.542; *p* < 0.05) but elevation did not (std *β* = −0.077; *p* = 0.70; Table 4). Populations with lower water temperatures displayed lower mean net immigration rates, indicating that they are the source populations (Fig. 4).

**Table 3.**
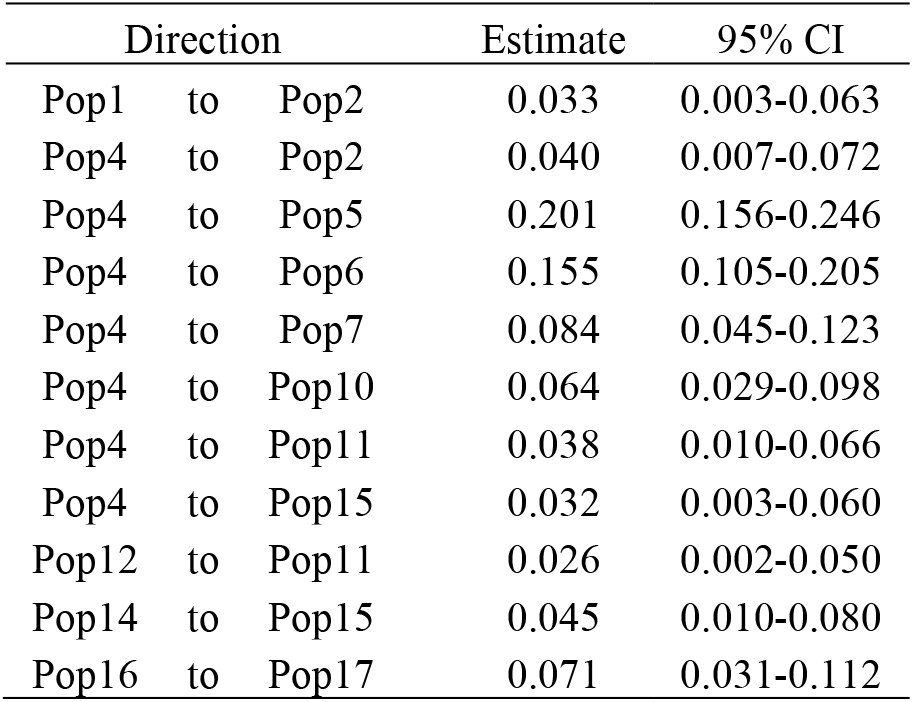
Significant gene flow estimated by BA3SNP (BayesAss). The estimated immigration rate and its 95% creditable interval (95% CI) are shown.

**Table 4.** The linear model explaining the mean net immigration rate. *P*-values were obtained by likelihood ratio tests.

| Variable | $\beta$ (SE) | std $\beta$ | $\chi^2$ | $p$ |
| --- | --- | --- | --- | --- |
| (Intercept) | -2.97*10<sup>-2</sup> (1.85*10<sup>-2</sup>) | -- | -- | -- |
| elevation | -7.86*10<sup>-6</sup> (2.21*10<sup>-5</sup>) | -0.077 | 0.15 | 0.70 |
| water temperature | 1.76*10<sup>-3</sup> (6.97*10<sup>-4</sup>) | 0.542 | 6.35 | <0.05 |

**Fig. 4.**
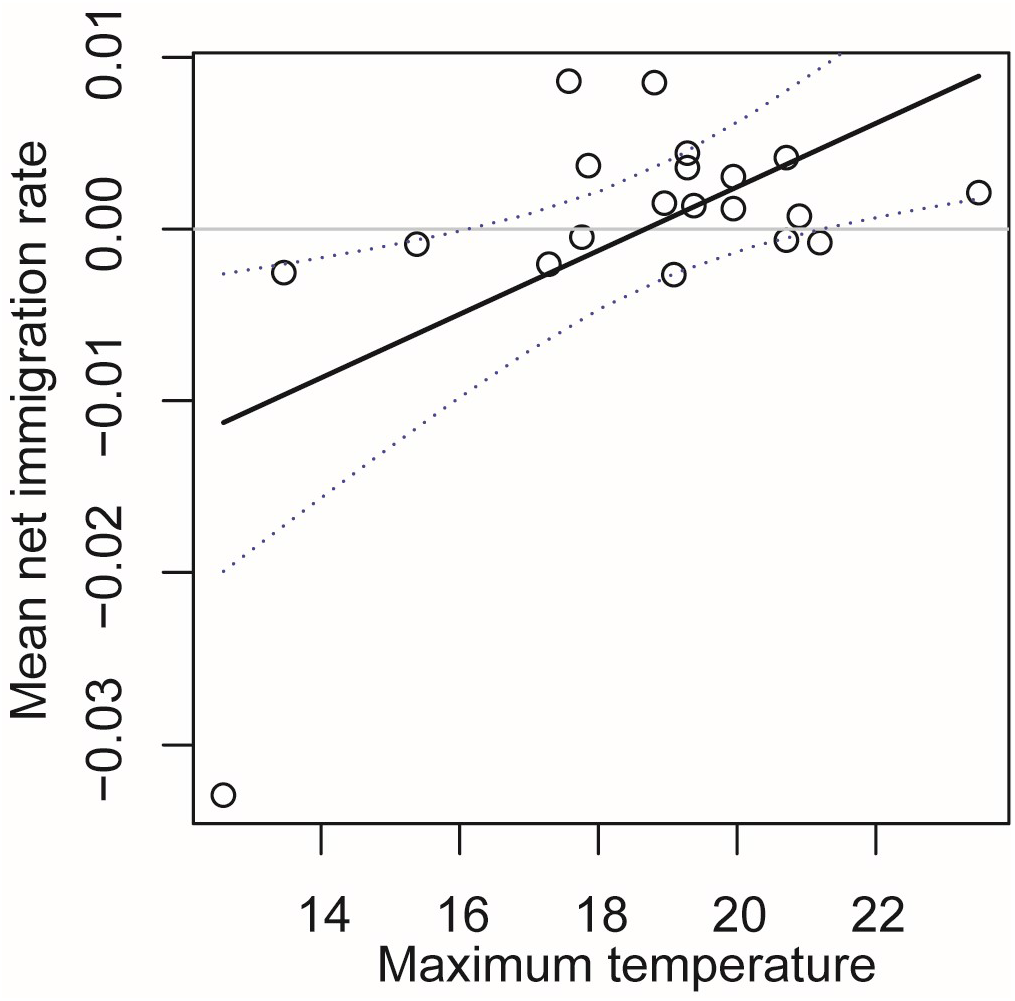
Variation in the mean net immigration rate with maximum water temperature. Populations with negative net immigration (i.e., greater emigration) indicate relative source populations, whereas populations with positive values (i.e., greater immigration) indicate relative sink populations.

## 4. Discussion

### 4.1. Genetic differentiation

Despite our expectation that the low mobility of *C. nozawae* would promote high genetic differentiation, a low level of genetic differentiation was observed (*F*_ST_ = 0.038). Other than gene flow, the large effective population size may result in low genetic drift from the common ancestor. Nevertheless, high-resolution genome-wide SNP analysis allowed genetic differentiation within the watershed to be clearly detected.

The genetic structure was distinctly defined within the watershed (Fig. 1). There was a significant correlation between genetic differentiation and geographic distance, but summer water temperature had no effect on the strength of genetic differentiation (Fig. 3). As the population density of *C. nozawae* has been shown to be largely determined by water temperature even within this thermal range, this result was contrary to our expectation that the signal of local adaptation can be found.

Only a handful of studies have associated the environment and genetic structure of sculpin. Some previous studies have reported that *Cottus* species display IBE patterns across different elevations or habitat types (Dennenmoser et al. 2014; Ruppert et al. 2017). However, these studies could not eliminate the interaction of environment and geographic distance. Of course, we cannot conclude that these studies in different localities and species have produced false positive results, but we should interpret these results carefully. In addition to the scarcity of previous studies, there will have been publication bias, with negative studies potentially not being published (Feder et al. 2012). Thus, the IBE of sculpin may be little in wild populations.

There may be some factors related to genetic differentiation that are not clear from this study (residuals). For example, the factors promoting the genetic divergence between Pop1-3 and Pop4-12 detected in STRUCTURE analysis are still unknown. Since we assume the IBE pattern as a hypothesis, we used only an environmental variable of each site, but the factor of genetic differentiation may also lie on the pathway between populations (e.g. streambed slope). To understand the residuals of genetic differentiation, further studies will be needed.

### 4.2. Gene flow

Another important insight into population structure was obtained via gene flow analysis. Almost all significant gene flow was detected in population pairs within approximately 15 km, the range where the spatial correlation was significant in the Mantel correlogram (Fig. 2B). Most of the 11 significant gene flow were downstream migration (Tables 3 and S2), which is consistent with several previous studies about fluvial sculpins that have revealed the source-sink structure from upstream to downstream sites (Hänfling and Weetman 2006; Junker et al. 2012). However, the present study also detected migrations toward tributaries at higher elevations, from lower-temperature tributaries to higher-temperature tributaries. Of course, we should be cautious about the accuracy of BayesAss estimates (Meirmans 2014). However, the estimated pattern displayed a clear trend, and it could be concluded that suitable habitats with low summer water temperatures behave as “source” habitats in the watershed. The effect of summer water temperature on the mean net immigration rate was larger than that of elevation (Table 4) and populations with lower temperatures had a greater degree of individual supply into other populations (Fig. 4).

In this study, significant gene flow was estimated between populations with different summer water temperatures at close locations (Fig. 1; Table 3). This type of gene flow, which in the direction of across dissimilar environments is high, is referred to as ‘counter-gradient gene flow’(Sexton et al. 2014). The IBE pattern is detected when gene flow among similar environments is high; therefore, the counter-gradient gene flow is the opposite pattern to IBE. Counter-gradient gene flow could result in migration load and therefore suppress adaptation due to migration (Sexton et al. 2014; Takahashi et al. 2016). The ecological processes driving this gene flow pattern could not be identified in this study, but it may at least be related to differences in population density (i.e., high density in low-temperature streams).

Goto (1998) used mark-recapture methods and reported that most mature individuals stay at the same site for more than one year, indicating almost no migration. While the mark-recapture method addresses only adult movement, genetic approaches also reflect movement during early life stages that is not easily captured by mark-recapture approaches (Lamphere and Blum 2012). If passive gene flow (i.e., the transportation of juveniles or eggs) is the driving force of gene flow, some high gene flow rates obtained in our study do not conflict with the results of mark-recapture methods. Furthermore, some previous studies on *Cottus* reported similar rates of gene flow to those obtained in the present study and discrepancies between genetic and mark-recapture estimates (Junker et al. 2012; Lamphere and Blum 2012), suggesting that the observed extent of gene flow is reasonable.

### 4.3 Implications for future management

Because sculpins are not commercially or recreationally important fish, they are impacted relatively little by human translocations (Lucek et al. 2018). Thus, sculpin genes have a footprint of a long-term past history, and the diversity of each watershed should be protected. Within the watershed, while each tributary is characterized by a unique genetic composition, loose connectivity via gene flow between tributaries is maintained. Indeed, fragmented populations such as Pop13 show lower genetic diversity. In particular, populations in streams with less suitable environments may be maintained by immigration from low-temperature streams that are more suitable habitats. Therefore, under ongoing climate change, (i) to preserve low-temperature streams and (ii) to prevent making unpassable barriers between source and sink populations would be the key measures for conserving sculpin populations. Considering that cold-water fish other than benthic fish have been well studied and revealed to show high mobility between tributaries, these measures will be effective for overall cold-water fish conservation. Groundwater and its associated factors such as watershed geology are useful for identifying low-water-temperature streams and conserving possible “source” populations.

Local conservation measures can also be proposed. If the distance at which the spatial coefficient becomes nonsignificant is considered to represent the appropriate management unit size (Diniz-Filho and De Campos Telles 2002), stream networks should not be subdivided into stretches of less than 15 km to maintain genetic resources. However, even if a large continuous habitat is maintained, then sections consisting solely of high-temperature streams will lead to local population declines due to the absence of individual supply sources. Thus, care must be taken whenever rivers are fragmented. In the Sorachi River, many *C. nozawae* inhabit high density (Suzuki, unpublished), and they are not considered to be in danger of immediate extinction. However, maintaining the abundance of general species is very important to avoid ecosystem breakdown (Baker et al. 2019). In some regions in Japan (Tohoku district), *C. nozawae* populations are in danger of local extinction (Ministry of the Environment Government of Japan 2020). Therefore, regardless of how large the population is, maintaining continuous stream networks and the connectivity to low-temperature sections in the networks is needed for future population persistence.

### 4.4 Implications for landscape/riverscape genetics

Decoupling distance and environment in a locality is a strong strategy for rigorously evaluating competing hypotheses (Gilbert and Lechowicz 2004). Through this approach, we revealed a pattern that has not been proposed for the examined genus. Freshwater populations have been identified as potentially fruitful targets for the application of a landscape genetic approach to delineate population structure (Kanno et al. 2011), and we focused on groundwater to reveal the genetic structure of cold-water fish populations. A groundwater-focused sampling design would be useful for purposes beyond revealing the effects of water temperature on population structure. The flow rate, which is another major factor in the stream environment, is also strongly influenced by groundwater. In addition, because groundwater flow is affected by not only geological conditions but also geomorphological conditions, it is possible to separate streams with high and low groundwater discharge even in the same geological watershed (e.g., Katahira et al. 2017), indicating that the elimination of correlations between distance and environment is also possible in smaller-scale studies.

In this study, the possibility of a source-sink structure caused by counter-gradient gene flow was suggested instead of IBE. Many source-sink structures from core populations to edge (peripheral) populations have been studied. However, local gene flow patterns are also important for discussing population structure in changing environments. The IBE pattern is easily detected when spatial-environmental correlation is present, but it is important to scrutinize whether it truly exists. Although the local source-sink pattern provides notable insights for conservation and evolution (Iles et al. 2018), a previous meta-analysis showed that counter-gradient gene flows are much less frequently reported than IBE (Sexton et al. 2014). Counter-gradient gene flow is inherently exclusive to IBE and may tend to be neglected. In population genetic studies, the results often differ from the hypotheses (Myers et al. 2019). As detecting the relationship between the environment and genetic differentiation is still challenging, we hope that the role of the environment in genetic divergence will be better understood in many studies with the help of sampling strategies.

## Acknowledgments

We are grateful to Jorge García Molinos for support with the data preparation. We thank the staff of the University of Tokyo Hokkaido Forest for their cooperation in selecting study sites. We also thank Suzuki K., Hotta W., Nishio T., Motosugi N., Kawai H., and Zakoh K. of Hokkaido University for their help in conducting the field sampling and laboratory work. This study is partly supported by the research fund for the Ishikari and Tokachi Rivers provided by the Ministry of Land, Infrastructure, Transport, and Tourism of Japan.

## Data Archiving

Raw MIG-seq reads were deposited in the DDBJ Sequence Read Archive under accession number DRA011249. The other data are available on Figshare (https://doi.org/10.6084/m9.figshare.13383245).

---

**Figure S1.** The values of posterior probability of the data (LnP(D)) from 20 runs for each value of K (left axis) and delta K (right axis) in the STRUCTURE analysis.

**Figure S2.** RDA biplots showing relationships between genetic data and MEMs generated by geographic distance. Significant MEM predictors are shown by the arrows. MEMs with smaller values model broad-scale spatial structures and MEMs with larger values model fine-scale spatial structures. Numbers in the plots represent sampling sites (population IDs). Adjusted R^2^ values (R_adj_^2^) and overall model significance are shown. (a) All 20 populations, Pop1-20; (b) Upstream 12 populations, Pop1-12; (c) Structured 9 populations, Pop4-12.

**Table S1.** Pairwise *F*_ST_ values between populations.

**Table S2.** Gene flow estimates obtained using BA3SNP (BayesAss). Column headings indicate the source populations, and row headings indicate the destination populations. The presented values represent the means of the posterior distributions of the migration rate into each population, and their standard deviation is provided in parentheses. Italics indicate the nonmigration rates, and boldface indicates significant gene flow.

## References

Almodóvar A, Nicola GG, Ayllón D, Elvira B (2012) Global warming threatens the persistence of Mediterranean brown trout. Glob Chang Biol 18: 1549–1560.

Arscott DB, Tockner K, Ward J V. (2001) Thermal heterogeneity along a braided floodplain river (Tagliamento River, northeastern Italy). Can J Fish Aquat Sci 58: 2359–2373.

Baker DJ, Garnett ST, O’Connor J, Ehmke G, Clarke RH, Woinarski JCZ, et al. (2019) Conserving the abundance of nonthreatened species. Conserv Biol 33: 319–328.

Brown LE, Milner AM, Hannah DM (2007) Groundwater influence on alpine stream ecosystems. Freshw Biol 52: 878–890.

Caissie D (2006) The thermal regime of rivers: A review. Freshw Biol 51: 1389–1406.

Catchen J, Hohenlohe PA, Bassham S, Amores A, Cresko WA (2013) Stacks: An analysis tool set for population genomics. Mol Ecol 22: 3124–3140.

Dennenmoser S, Rogers SM, Vamosi SM (2014) Genetic population structure in prickly sculpin (Cottus asper) reflects isolation-by-environment between two life-history ecotypes. Biol J Linn Soc 113: 943–957.

Diniz-Filho JAF, De Campos Telles MP (2002) Spatial autocorrelation analysis and the identification of operational units for conservation in continuous populations. Conserv Biol 16: 924–935.

Diniz-Filho JAF, Soares TN, Lima JS, Dobrovolski R, Landeiro VL, Telles MP de C, et al. (2013) Mantel test in population genetics. Genet Mol Biol 36: 475–485.

Dray S, Bauman D, Blanchet G, Borcard D, Clappe S, Guenard G, et al. (2020) adespatial: Multivariate Multiscale Spatial Analysis. R package version 0.3-8. https://CRAN.R-project.org/package=adespatial

Earl DA, vonHoldt BM (2012) STRUCTURE HARVESTER: A website and program for visualizing STRUCTURE output and implementing the Evanno method. Conserv Genet Resour 4: 359–361.

Evanno G, Regnaut S, Goudet J (2005) Detecting the number of clusters of individuals using the software STRUCTURE: A simulation study. Mol Ecol 14: 2611–2620.

Feder JL, Egan SP, Nosil P (2012) The genomics of speciation-with-gene-flow. Trends Genet 28: 342–350.

Gilbert B, Bennett JR (2010) Partitioning variation in ecological communities: Do the numbers add up? J Appl Ecol 47: 1071–1082.

Gilbert B, Lechowicz MJ (2004) Neutrality, niches, and dispersal in a temperate forest understory. Proc Natl Acad Sci U S A 101: 7651–7656.

Goslee S, Urban D (2007) The ecodist package for dissimilarity-based analysis of ecological data. J Stat Softw 22: 1–19.

Goto A (1998) Life-history variations in the fluvial sculpin, Cottus nozawae (Cottidae), along the course of a small mountain stream. Environ Biol Fishes 52: 203–212.

Goudet J (1995) FSTAT (Version 1.2): A Computer Program to Calculate F-Statistics. J Hered 86: 485–486.

Griffith DA, Peres-Neto PR (2006) Spatial modeling in ecology: The flexibility of eigenfunction spatial analyses. Ecology 87: 2603–2613.

Guillot G, Rousset F (2013) Dismantling the Mantel tests. Methods Ecol Evol 4: 336–344.

Hänfling B, Weetman D (2006) Concordant genetic estimators of migration reveal anthropogenically enhanced source-sink population structure in the river sculpin, Cottus gobio. Genetics 173: 1487–1501.

Harmon LJ, Glor RE (2010) Poor statistical performance of the mantel test in phylogenetic comparative analyses. Evolution 64: 2173–2178.

Heino J, Grönroos M, Ilmonen J, Karhu T, Niva M, Paasivirta L (2013) Environmental heterogeneity and β diversity of stream macroinvertebrate communities at intermediate spatial scales. Freshw Sci 32: 142–154.

Hohenlohe PA, Funk WC, Rajora OP (2021) Population genomics for wildlife conservation and management. Mol Ecol 30: 62–82

Hubisz MJ, Falush D, Stephens M, Pritchard JK (2009) Inferring weak population structure with the assistance of sample group information. Mol Ecol Resour 9: 1322–1332.

Iles DT, Williams NM, Crone EE (2018) Source-sink dynamics of bumblebees in rapidly changing landscapes. J Appl Ecol 55: 2802–2811.

Junker J, Peter A, Wagner CE, Mwaiko S, Germann B, Seehausen O, et al. (2012) River fragmentation increases localized population genetic structure and enhances asymmetry of dispersal in bullhead (Cottus gobio). Conserv Genet 13: 545–556.

Kanno Y, Vokoun JC, Letcher BH (2011) Fine-scale population structure and riverscape genetics of brook trout (Salvelinus fontinalis) distributed continuously along headwater channel networks. Mol Ecol 20: 3711–3729.

Katahira H, Yamazaki C, Fukui S, Ayer CG, Koizumi I (2017) Spatial aggregation in small spring-fed tributaries leads to a potential metapopulation structure in a parasitic fish leech. Parasitol Open 3.

Kawecki TJ, Ebert D (2004) Conceptual issues in local adaptation. Ecol Lett 7: 1225–1241.

Koizumi I (2011) Integration of ecology, demography and genetics to reveal population structure and persistence: A mini review and case study of stream-dwelling Dolly Varden. Ecol Freshw Fish 20: 352–363.

Koizumi I, Maekawa K (2004) Metapopulation structure of stream-dwelling Dolly Varden charr inferred from patterns of occurrence in the Sorachi River basin, Hokkaido, Japan. Freshw Biol 49: 973–981.

Koizumi I, Yamamoto S, Maekawa K (2006) Decomposed pairwise regression analysis of genetic and geographic distances reveals a metapopulation structure of stream-dwelling Dolly Varden charr. Mol Ecol 15: 3175–3189.

Kopelman NM, Mayzel J, Jakobsson M, Rosenberg NA, Mayrose I (2015) Clumpak: A program for identifying clustering modes and packaging population structure inferences across K. Mol Ecol Resour 15: 1179–1191.

Lamphere BA, Blum MJ (2012) Genetic estimates of population structure and dispersal in a benthic stream fish. Ecol Freshw Fish 21: 75–86.

Legendre P, Fortin MJ, Borcard D (2015) Should the Mantel test be used in spatial analysis? Methods Ecol Evol 6: 1239–1247.

Legendre P, Legendre L (2012) Numerical Ecology.

Lichstein JW (2007) Multiple regression on distance matrices: A multivariate spatial analysis tool. Plant Ecol 188: 117–131.

Lucek K, Keller I, Nolte AW, Seehausen O (2018) Distinct colonization waves underlie the diversification of the freshwater sculpin (Cottus gobio) in the Central European Alpine region. J Evol Biol 31: 1254–1267.

Meirmans PG (2012) The trouble with isolation by distance. Mol Ecol 21: 2839–2846.

Meirmans PG (2014) Nonconvergence in Bayesian estimation of migration rates. Mol Ecol Resour 14: 726–733.

Meirmans PG (2015) Seven common mistakes in population genetics and how to avoid them. Mol Ecol 24: 3223–3231.

Middaugh CR, Kessinger B, Magoulick DD (2018) Climate-induced seasonal changes in smallmouth bass growth rate potential at the southern range extent. Ecol Freshw Fish 27: 19–29.

Ministry of the Environment Government of Japan (2020) Red List of Japan.

Mussmann SM, Douglas MR, Chafin TK, Douglas ME (2019) BA3-SNPs: Contemporary migration reconfigured in BayesAss for next-generation sequence data. Methods Ecol Evol 10: 1808–1813.

Myers EA, Xue AT, Gehara M, Cox CL, Davis Rabosky AR, Lemos-Espinal J, et al. (2019) Environmental heterogeneity and not vicariant biogeographic barriers generate community-wide population structure in desert-adapted snakes. Mol Ecol 28: 4535–4548.

Nagasaka A, Sugiyama S (2010) Factors affecting the summer maximum stream temperature of small streams in northern Japan. Bull Hokkaido For Res Inst 47: 35–43. (In Japanese with English abstract)

Nakajima S, Hirota SK, Matsuo A, Suyama Y, Nakamura F (2020) Genetic structure and population demography of white-spotted charr in the upstream watershed of a large dam. Water 12: 2406.

Nakamura F, Yamada H (2005) Effects of pasture development on the ecological functions of riparian forests in Hokkaido in northern Japan. Ecol Eng 24: 539–550.

Nosil P, Vines TH, Funk DJ (2005) Perspective: reproductive isolation caused by natural selection against immigrants from divergent habitats. Evolution 59: 705–719.

Oksanen JF, Blanchet G, Friendly M, Kindt R, Legendre P, McGlinn D, et al. (2019) vegan: Community Ecology Package. R package version 2.5-6. https://CRAN.R-project.org/package=vegan

Okumura N, Goto A (1996) Genetic variation and differentiation of the two river sculpins, Cottus nozawae and C. amblystomopsis, deduced from allozyme and restriction enzyme-digested mtDNA fragment length polymorphism analyses. Ichthyol Res 43: 399–416.

Olden JD, Naiman RJ (2010) Incorporating thermal regimes into environmental flows assessments: Modifying dam operations to restore freshwater ecosystem integrity. Freshw Biol 55: 86–107.

Orsini L, Vanoverbeke J, Swillen I, Mergeay J, De Meester L (2013) Drivers of population genetic differentiation in the wild: Isolation by dispersal limitation, isolation by adaptation and isolation by colonization. Mol Ecol 22: 5983–5999.

Peakall R, Smouse PE (2012) GenAlEx 6.5: Genetic analysis in Excel. Population genetic software for teaching and research-an update. Bioinformatics 28: 2537–2539.

Poff NL, Richter BD, Arthington AH, Bunn SE, Naiman RJ, Kendy E, et al. (2010) The ecological limits of hydrologic alteration (ELOHA): A new framework for developing regional environmental flow standards. Freshw Biol 55: 147–170.

Pritchard JK, Stephens M, Donnelly P (2000) Inference of Population Structure Using Multilocus Genotype Data. Genetics 155: 945–959.

Raufaste N, Francois R (2001) Are Partial Mantel Test Adequate? Evolution 55: 1703–1705.

Richardson JL, Brady SP, Wang IJ, Spear SF (2016) Navigating the pitfalls and promise of landscape genetics. Mol Ecol 25: 849–864.

R Core Team (2019) R: A language and environment for statistical computing. R Foundation for Statistical Computing, Vienna, Austria. https://www.R-project.org/.

Rousset F (2002) Partial Mantel tests: Reply to Castellano and Balletto. Evolution 56: 1874–1875.

Ruppert JLW, James PMA, Taylor EB, Rudolfsen T, Veillard M, Davis CS, et al. (2017) Riverscape genetic structure of a threatened and dispersal limited freshwater species, the Rocky Mountain Sculpin (Cottus sp.). Conserv Genet 18: 925–937.

Sexton JP, Hangartner SB, Hoffmann AA (2014) Genetic isolation by environment or distance: Which pattern of gene flow is most common? Evolution 68: 1–15.

Sexton JP, Hufford MB, Bateman AC, Lowry DB, Meimberg H, Strauss SY, et al. (2016) Climate structures genetic variation across a species’ elevation range: A test of range limits hypotheses. Mol Ecol 25: 911–928.

Suyama Y, Matsuki Y (2015) MIG-seq: An effective PCR-based method for genome-wide single-nucleotide polymorphism genotyping using the next-generation sequencing platform. Sci Rep 5: 1–12.

Tague CL, Farrell M, Grant G, Lewis S, Rey S (2007) Hydrogeologic controls on summer stream temperatures in the McKenzie River basin, Oregon. Hydrol Process 21: 3288–3300.

Takahashi Y, Suyama Y, Matsuki Y, Funayama R, Nakayama K, Kawata M (2016) Lack of genetic variation prevents adaptation at the geographic range margin in a damselfly. Mol Ecol 25: 4450–4460.

Thorpe RS, Surget-Groba Y, Johansson H (2008) The relative importance of ecology and geographic isolation for speciation in anoles. Philos Trans R Soc B Biol Sci 363: 3071–3081.

Tsuda Y, Nakao K, Ide Y, Tsumura Y (2015) The population demography of Betula maximowicziana, a cool-temperate tree species in Japan, in relation to the last glacial period: Its admixture-like genetic structure is the result of simple population splitting not admixing. Mol Ecol 24: 1403–1418.

Uno H (2016) Stream thermal heterogeneity prolongs aquatic-terrestrial subsidy and enhances riparian spider growth. Ecology 97: 2547–2553.

Wang IJ, Bradburd GS (2014) Isolation by environment. Mol Ecol 23: 5649–5662.

Wang IJ, Summers K (2010) Genetic structure is correlated with phenotypic divergence rather than geographic isolation in the highly polymorphic strawberry poison-dart frog. Mol Ecol 19: 447–458.

Watz J, Otsuki Y, Nagatsuka K, Hasegawa K, Koizumi I (2019) Temperature-dependent competition between juvenile salmonids in small streams. Freshw Biol 64: 1534–1541.

Weir BS, Cockerham CC (1984) Estimating F-statistics for the analysis of population structure. Evolution 38: 1358–1370

Wilson GA, Rannala B (2003) Bayesian inference of recent migration rates using multilocus genotypes. Genetics 163: 1177–1191.

Wright S (1943) Isolation by Distance. Genetics 28: 114–38.

Yagami T, Goto A (2000) Patchy distribution of a fluvial sculpin, Cottus nozawae, in the Gakko River system at the southern margin of its native range. Ichthyol Res 47: 277–286.

Zeileis A, Hothorn T (2002) Diagnostic checking in regression relationships. R News 2: 7–10. https://CRAN.R-project.org/doc/Rnews/.

Zeller KA, Creech TG, Millette KL, Crowhurst RS, Long RA, Wagner HH, et al. (2016) Using simulations to evaluate Mantel-based methods for assessing landscape resistance to gene flow. Ecol Evol 6: 4115–4128.

