## Supplemental information for "A strategic sampling design revealed the local genetic structure of cold-water fluvial sculpin: a focus on groundwater-dependent water temperature heterogeneity"

**Figure S1.** The values of posterior probability of the data (LnP(D)) from 20 runs for each value of K (left axis) and delta K (right axis) in the STRUCTURE analysis.

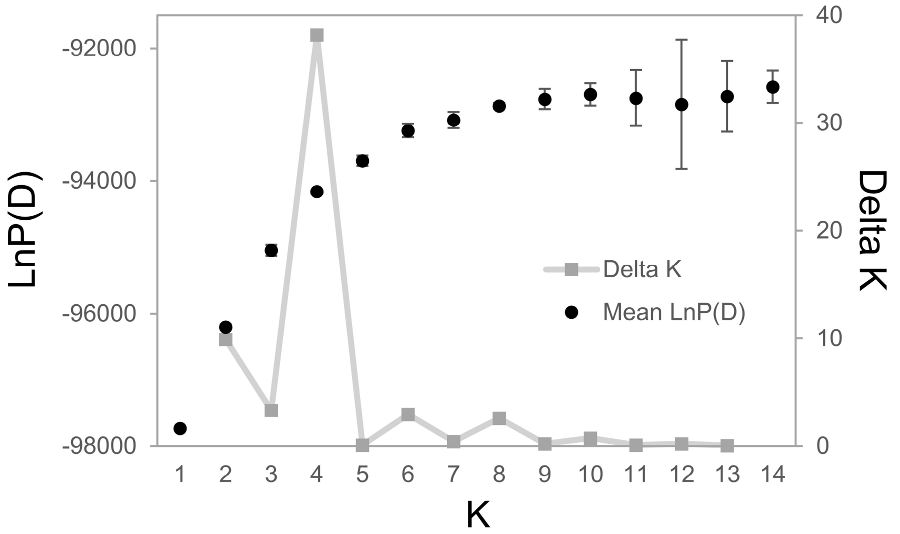

**Figure S2.** RDA biplots showing relationships between genetic data and MEMs generated by geographic distance. Significant MEM predictors are shown by the arrows. MEMs with smaller values model broad-scale spatial structures and MEMs with larger values model fine-scale spatial structures. Numbers in the plots represent sampling sites (population IDs). Adjusted $R^2$ values ($R_{adj}^2$) and overall model significance are shown. (a) All 20 populations, Pop1-20; (b) Upstream 12 populations, Pop1-12; (c) Structured 9 populations, Pop4-12.

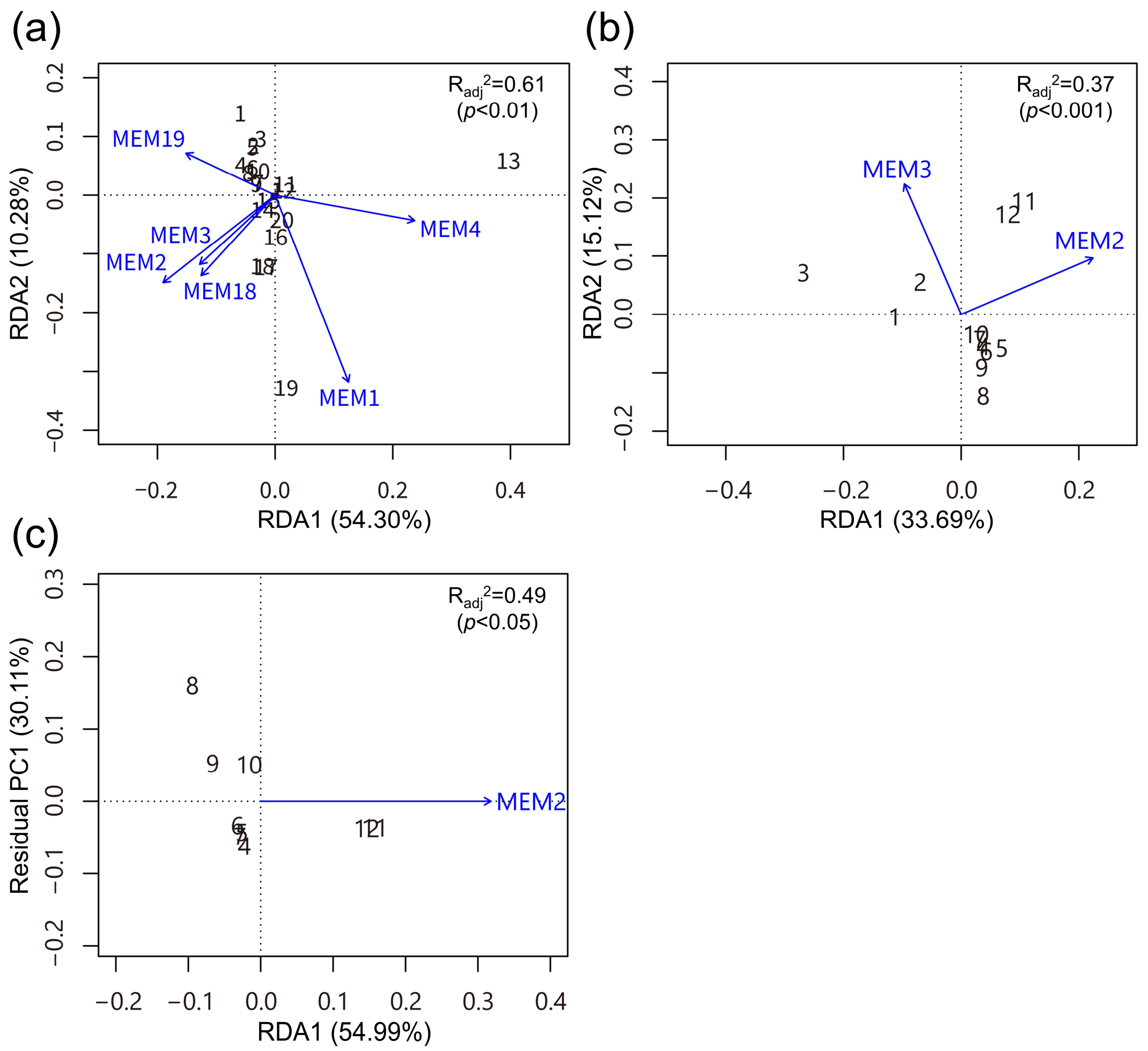

**Table S1.** Pairwise $F_{ST}$ values between populations.

| | Pop1 | Pop2 | Pop3 | Pop4 | Pop5 | Pop6 | Pop7 | Pop8 | Pop9 | Pop10 | Pop11 | Pop12 | Pop13 | Pop14 | Pop15 | Pop16 | Pop17 | Pop18 | Pop19 | Pop20 |
|---|---|---|---|---|---|---|---|---|---|---|---|---|---|---|---|---|---|---|---|---|
| Pop1 | 0.000 | | | | | | | | | | | | | | | | | | | |
| Pop2 | 0.021 | 0.000 | | | | | | | | | | | | | | | | | | |
| Pop3 | 0.050 | 0.041 | 0.000 | | | | | | | | | | | | | | | | | |
| Pop4 | 0.034 | 0.025 | 0.049 | 0.000 | | | | | | | | | | | | | | | | |
| Pop5 | 0.038 | 0.029 | 0.056 | 0.001 | 0.000 | | | | | | | | | | | | | | | |
| Pop6 | 0.037 | 0.026 | 0.050 | 0.004 | 0.002 | 0.000 | | | | | | | | | | | | | | |
| Pop7 | 0.029 | 0.022 | 0.046 | 0.006 | 0.007 | 0.006 | 0.000 | | | | | | | | | | | | | |
| Pop8 | 0.039 | 0.037 | 0.054 | 0.019 | 0.014 | 0.012 | 0.005 | 0.000 | | | | | | | | | | | | |
| Pop9 | 0.041 | 0.030 | 0.055 | 0.020 | 0.017 | 0.016 | 0.008 | 0.004 | 0.000 | | | | | | | | | | | |
| Pop10 | 0.030 | 0.020 | 0.043 | 0.009 | 0.007 | 0.004 | 0.004 | 0.006 | 0.004 | 0.000 | | | | | | | | | | |
| Pop11 | 0.051 | 0.034 | 0.065 | 0.028 | 0.027 | 0.028 | 0.024 | 0.040 | 0.035 | 0.023 | 0.000 | | | | | | | | | |
| Pop12 | 0.050 | 0.034 | 0.058 | 0.026 | 0.024 | 0.026 | 0.024 | 0.038 | 0.035 | 0.021 | 0.007 | 0.000 | | | | | | | | |
| Pop13 | 0.127 | 0.109 | 0.125 | 0.116 | 0.108 | 0.105 | 0.098 | 0.114 | 0.105 | 0.098 | 0.088 | 0.085 | 0.000 | | | | | | | |
| Pop14 | 0.040 | 0.028 | 0.051 | 0.021 | 0.032 | 0.029 | 0.027 | 0.042 | 0.033 | 0.026 | 0.040 | 0.032 | 0.104 | 0.000 | | | | | | |
| Pop15 | 0.038 | 0.027 | 0.049 | 0.022 | 0.028 | 0.029 | 0.025 | 0.039 | 0.029 | 0.024 | 0.033 | 0.028 | 0.096 | 0.006 | 0.000 | | | | | |
| Pop16 | 0.033 | 0.033 | 0.043 | 0.033 | 0.035 | 0.032 | 0.027 | 0.043 | 0.039 | 0.031 | 0.037 | 0.035 | 0.091 | 0.022 | 0.023 | 0.000 | | | | |
| Pop17 | 0.035 | 0.031 | 0.045 | 0.028 | 0.036 | 0.030 | 0.026 | 0.038 | 0.036 | 0.026 | 0.037 | 0.032 | 0.099 | 0.016 | 0.020 | 0.006 | 0.000 | | | |
| Pop18 | 0.039 | 0.035 | 0.051 | 0.027 | 0.034 | 0.024 | 0.022 | 0.025 | 0.021 | 0.023 | 0.044 | 0.037 | 0.102 | 0.025 | 0.022 | 0.010 | 0.011 | 0.000 | | |
| Pop19 | 0.081 | 0.063 | 0.080 | 0.059 | 0.058 | 0.053 | 0.043 | 0.051 | 0.047 | 0.047 | 0.050 | 0.049 | 0.115 | 0.059 | 0.057 | 0.051 | 0.038 | 0.034 | 0.000 | |
| Pop20 | 0.055 | 0.038 | 0.061 | 0.029 | 0.030 | 0.028 | 0.022 | 0.028 | 0.022 | 0.022 | 0.021 | 0.023 | 0.087 | 0.025 | 0.017 | 0.036 | 0.034 | 0.031 | 0.042 | 0.000 |

**Table S2.** Gene flow estimates obtained using BA3SNP (BayesAss). Column headings indicate the source populations, and row headings indicate the destination populations. The presented values represent the means of the posterior distributions of the migration rate into each population, and their standard deviation is provided in parentheses. Italics indicate the nonmigration rates, and boldface indicates significant gene flow.

| | <-Pop1 | <-Pop2 | <-Pop3 | <-Pop4 | <-Pop5 | <-Pop6 | <-Pop7 | <-Pop8 | <-Pop9 | <-Pop10 | <-Pop11 | <-Pop12 | <-Pop13 | <-Pop14 | <-Pop15 | <-Pop16 | <-Pop17 | <-Pop18 | <-Pop19 | <-Pop20 |
|---|---|---|---|---|---|---|---|---|---|---|---|---|---|---|---|---|---|---|---|---|
| Pop1 | *0.8724(0.0223)* | 0.0062(0.0061) | 0.0064(0.0064) | 0.0064(0.0062) | 0.0067(0.0065) | 0.0064(0.0063) | 0.0064(0.0064) | 0.0062(0.0060) | 0.0065(0.0062) | 0.0063(0.0064) | 0.0063(0.0063) | 0.0063(0.0061) | 0.0062(0.0061) | 0.0065(0.0064) | 0.0064(0.0062) | 0.0064(0.0063) | 0.0062(0.0063) | 0.0128(0.0088) | 0.0065(0.0063) | 0.0063(0.0062) |
| Pop2 | **0.0331(0.0153)** | *0.7920(0.0244)* | 0.0078(0.0077) | **0.0397(0.0167)** | 0.0081(0.0080) | 0.0078(0.0075) | 0.0081(0.0079) | 0.0080(0.0077) | 0.0079(0.0077) | 0.0080(0.0082) | 0.0080(0.0079) | 0.0078(0.0079) | 0.0080(0.0077) | 0.0079(0.0077) | 0.0079(0.0075) | 0.0080(0.0078) | 0.0079(0.0079) | 0.0081(0.0077) | 0.0081(0.0076) | 0.0080(0.0078) |
| Pop3 | 0.0065(0.0065) | 0.0130(0.0091) | *0.8695(0.0224)* | 0.0064(0.0063) | 0.0066(0.0064) | 0.0065(0.0065) | 0.0067(0.0068) | 0.0064(0.0063) | 0.0066(0.0066) | 0.0066(0.0065) | 0.0066(0.0066) | 0.0064(0.0063) | 0.0064(0.0061) | 0.0066(0.0064) | 0.0067(0.0063) | 0.0064(0.0062) | 0.0064(0.0063) | 0.0063(0.0062) | 0.0066(0.0066) | 0.0066(0.0065) |
| Pop4 | 0.0060(0.0060) | 0.0064(0.0061) | 0.0061(0.0059) | *0.8827(0.0216)* | 0.0063(0.0061) | 0.0063(0.0062) | 0.0061(0.0061) | 0.0063(0.0061) | 0.0062(0.0060) | 0.0062(0.0062) | 0.0061(0.0060) | 0.0060(0.0059) | 0.0062(0.0062) | 0.0063(0.0061) | 0.0062(0.0062) | 0.0063(0.0064) | 0.0062(0.0060) | 0.0060(0.0058) | 0.0062(0.0060) | 0.0061(0.0060) |
| Pop5 | 0.0064(0.0062) | 0.0066(0.0065) | 0.0068(0.0067) | **0.2007(0.0230)** | *0.6802(0.0091)* | 0.0067(0.0064) | 0.0065(0.0063) | 0.0064(0.0063) | 0.0066(0.0065) | 0.0069(0.0067) | 0.0066(0.0064) | 0.0066(0.0065) | 0.0067(0.0066) | 0.0066(0.0066) | 0.0068(0.0067) | 0.0065(0.0065) | 0.0066(0.0065) | 0.0065(0.0065) | 0.0067(0.0067) | 0.0066(0.0065) |
| Pop6 | 0.0080(0.0079) | 0.0081(0.0079) | 0.0081(0.0079) | **0.1549(0.0255)** | 0.0084(0.0081) | *0.6993(0.0153)* | 0.0080(0.0078) | 0.0080(0.0078) | 0.0082(0.0080) | 0.0084(0.0080) | 0.0078(0.0076) | 0.0081(0.0081) | 0.0081(0.0078) | 0.0080(0.0077) | 0.0083(0.0082) | 0.0081(0.0080) | 0.0080(0.0079) | 0.0080(0.0076) | 0.0079(0.0075) | 0.0082(0.0077) |
| Pop7 | 0.0126(0.0087) | 0.0129(0.0089) | 0.0064(0.0063) | **0.0838(0.0200)** | 0.0125(0.0085) | 0.0064(0.0065) | *0.7575(0.0206)* | 0.0063(0.0062) | 0.0128(0.0088) | 0.0131(0.0091) | 0.0064(0.0063) | 0.0062(0.0064) | 0.0065(0.0064) | 0.0182(0.0107) | 0.0065(0.0064) | 0.0062(0.0062) | 0.0062(0.0062) | 0.0064(0.0062) | 0.0066(0.0066) | 0.0065(0.0066) |
| Pop8 | 0.0062(0.0060) | 0.0063(0.0063) | 0.0065(0.0063) | 0.0128(0.0087) | 0.0064(0.0062) | 0.0062(0.0062) | 0.0128(0.0086) | *0.8529(0.0230)* | 0.0192(0.0108) | 0.0064(0.0064) | 0.0064(0.0062) | 0.0063(0.0062) | 0.0065(0.0065) | 0.0066(0.0065) | 0.0062(0.0062) | 0.0065(0.0062) | 0.0063(0.0062) | 0.0066(0.0065) | 0.0065(0.0062) | 0.0065(0.0065) |
| Pop9 | 0.0066(0.0066) | 0.0063(0.0063) | 0.0063(0.0061) | 0.0129(0.0090) | 0.0063(0.0063) | 0.0062(0.0061) | 0.0126(0.0086) | 0.0066(0.0065) | *0.8592(0.0230)* | 0.0129(0.0091) | 0.0063(0.0063) | 0.0066(0.0064) | 0.0065(0.0063) | 0.0063(0.0062) | 0.0065(0.0064) | 0.0064(0.0062) | 0.0061(0.0061) | 0.0063(0.0064) | 0.0064(0.0064) | 0.0065(0.0063) |
| Pop10 | 0.0065(0.0065) | 0.0063(0.0062) | 0.0063(0.0061) | **0.0638(0.0177)** | 0.0191(0.0107) | 0.0064(0.0064) | 0.0128(0.0088) | 0.0195(0.0109) | 0.0191(0.0106) | *0.7629(0.0206)* | 0.0129(0.0090) | 0.0128(0.0088) | 0.0065(0.0064) | 0.0064(0.0062) | 0.0063(0.0062) | 0.0064(0.0063) | 0.0064(0.0064) | 0.0065(0.0063) | 0.0065(0.0065) | 0.0065(0.0063) |
| Pop11 | 0.0065(0.0063) | 0.0065(0.0063) | 0.0065(0.0065) | **0.0381(0.0144)** | 0.0064(0.0063) | 0.0065(0.0062) | 0.0062(0.0061) | 0.0063(0.0063) | 0.0064(0.0063) | 0.0066(0.0065) | *0.8266(0.0227)* | **0.0258(0.0123)** | 0.0064(0.0064) | 0.0063(0.0064) | 0.0065(0.0063) | 0.0064(0.0064) | 0.0062(0.0062) | 0.0066(0.0065) | 0.0066(0.0063) | 0.0064(0.0064) |
| Pop12 | 0.0064(0.0062) | 0.0063(0.0061) | 0.0063(0.0060) | 0.0188(0.0103) | 0.0065(0.0063) | 0.0065(0.0064) | 0.0063(0.0063) | 0.0064(0.0061) | 0.0063(0.0062) | 0.0065(0.0063) | 0.0129(0.0088) | *0.8535(0.0228)* | 0.0061(0.0060) | 0.0064(0.0063) | 0.0064(0.0062) | 0.0130(0.0091) | 0.0063(0.0062) | 0.0062(0.0062) | 0.0064(0.0063) | 0.0066(0.0065) |
| Pop13 | 0.0079(0.0077) | 0.0081(0.0079) | 0.0083(0.0080) | 0.0079(0.0076) | 0.0080(0.0076) | 0.0082(0.0079) | 0.0081(0.0078) | 0.0081(0.0077) | 0.0079(0.0075) | 0.0079(0.0078) | 0.0079(0.0078) | 0.0078(0.0076) | *0.8489(0.0251)* | 0.0079(0.0076) | 0.0078(0.0074) | 0.0077(0.0074) | 0.0079(0.0078) | 0.0081(0.0078) | 0.0077(0.0074) | 0.0080(0.0079) |
| Pop14 | 0.0081(0.0077) | 0.0083(0.0081) | 0.0081(0.0079) | 0.0083(0.0079) | 0.0081(0.0077) | 0.0083(0.0082) | 0.0084(0.0083) | 0.0079(0.0078) | 0.0081(0.0080) | 0.0083(0.0080) | 0.0079(0.0078) | 0.0080(0.0081) | 0.0082(0.0079) | *0.8287(0.0260)* | 0.0247(0.0136) | 0.0083(0.0080) | 0.0079(0.0077) | 0.0082(0.0081) | 0.0081(0.0082) | 0.0080(0.0078) |
| Pop15 | 0.0076(0.0075) | 0.0079(0.0077) | 0.0078(0.0077) | **0.0315(0.0147)** | 0.0080(0.0079) | 0.0081(0.0078) | 0.0077(0.0074) | 0.0078(0.0076) | 0.0080(0.0078) | 0.0080(0.0078) | 0.0078(0.0075) | 0.0079(0.0077) | 0.0080(0.0076) | **0.0451(0.0180)** | *0.7894(0.0248)* | 0.0079(0.0078) | 0.0077(0.0074) | 0.0080(0.0079) | 0.0077(0.0077) | 0.0080(0.0078) |
| Pop16 | 0.0079(0.0079) | 0.0078(0.0077) | 0.0083(0.0083) | 0.0157(0.0108) | 0.0079(0.0078) | 0.0081(0.0077) | 0.0079(0.0077) | 0.0079(0.0077) | 0.0079(0.0077) | 0.0079(0.0077) | 0.0077(0.0077) | 0.0079(0.0077) | 0.0080(0.0079) | 0.0079(0.0076) | 0.0079(0.0079) | *0.8414(0.0258)* | 0.0082(0.0078) | 0.0078(0.0077) | 0.0079(0.0078) | 0.0080(0.0076) |
| Pop17 | 0.0081(0.0077) | 0.0079(0.0078) | 0.0081(0.0078) | 0.0079(0.0076) | 0.0078(0.0074) | 0.0077(0.0078) | 0.0079(0.0078) | 0.0081(0.0078) | 0.0081(0.0079) | 0.0080(0.0079) | 0.0078(0.0076) | 0.0079(0.0077) | 0.0076(0.0073) | 0.0079(0.0076) | 0.0078(0.0077) | **0.0714(0.0208)** | *0.7859(0.0245)* | 0.0080(0.0080) | 0.0081(0.0078) | 0.0079(0.0076) |
| Pop18 | 0.0087(0.0085) | 0.0085(0.0080) | 0.0086(0.0084) | 0.0085(0.0085) | 0.0086(0.0085) | 0.0083(0.0081) | 0.0086(0.0082) | 0.0168(0.0112) | 0.0084(0.0080) | 0.0086(0.0082) | 0.0085(0.0083) | 0.0086(0.0083) | 0.0088(0.0085) | 0.0089(0.0084) | 0.0085(0.0082) | 0.0085(0.0083) | 0.0086(0.0084) | *0.8286(0.0261)* | 0.0087(0.0084) | 0.0087(0.0083) |
| Pop19 | 0.0082(0.0081) | 0.0082(0.0079) | 0.0082(0.0078) | 0.0084(0.0079) | 0.0079(0.0078) | 0.0081(0.0078) | 0.0083(0.0079) | 0.0080(0.0079) | 0.0081(0.0079) | 0.0082(0.0079) | 0.0081(0.0080) | 0.0081(0.0080) | 0.0082(0.0078) | 0.0082(0.0078) | 0.0082(0.0081) | 0.0081(0.0080) | 0.0164(0.0113) | 0.0080(0.0077) | *0.8371(0.0259)* | 0.0081(0.0079) |
| Pop20 | 0.0168(0.0116) | 0.0084(0.0082) | 0.0083(0.0083) | 0.0165(0.0113) | 0.0084(0.0082) | 0.0084(0.0080) | 0.0249(0.0136) | 0.0084(0.0081) | 0.0170(0.0114) | 0.0082(0.0080) | 0.0084(0.0081) | 0.0081(0.0078) | 0.0083(0.0083) | 0.0085(0.0082) | 0.0247(0.0136) | 0.0082(0.0082) | 0.0085(0.0083) | 0.0086(0.0084) | 0.0079(0.0080) | *0.7834(0.0253)* |
